# New Insights into Neuromuscular Contracture Reveals Myotendinous-SMAD4 Signaling Underlies Contracture Formation

**DOI:** 10.1101/2024.06.11.598573

**Authors:** Varun Arvind, Peter Timothy Shyu, Joshua E. Hyman, Alice H. Huang

**Affiliations:** Columbia University

## Abstract

Neuromuscular contractures (NC) are a prevalent cause of joint deformity in children suffering from neuromuscular disorders or nerve damage, leading to persistent disability. The role of tendon in the development of NC remains poorly understood, with current treatments predominantly targeting muscle. Here, we establish a surgical model of NC in the hindlimb that recapitulates functional deformity and transcriptomic changes observed in human disease. Our findings indicate that in NC, tendons dramatically elongate, undergoing changes in matrix and structural composition that reduce tensile stiffness. Contrary to expectations, we find that tendon elongation was principally driven by increased myotendon infiltration into muscle which restricted muscle elongation contributing to NC. Using lineage tracing, we show that myotendon elongation was due to increased infiltration of intrinsically derived tenocytes. Transcriptional profiling revealed BMP signaling as a key factor in myotendon elongation, corroborated by elevated myotendinous Smad4 activity in both our mouse model and in human NC tissues. Crucially, administration of a small molecule inhibitor of BMP-mediated Smad4 signaling not only restored joint mobility but also prevented myotendon elongation. These insights establish of a clinically relevant mouse model of NC and unveil a novel role for myotendon elongation in NC progression. Excitingly, our results suggest that targeting myotendon signaling could represent a new direction for tendon-focused therapies in NC management.

## INTRODUCTION

A principal role of the musculoskeletal system is to provide stability while also accommodating a high range of motion across joints. During development, precise organization of the muscle-tendon attachment to bone creates fixed anchor points that transmit forces from muscle to bone and permit joint motion(Felsenthal et al., 2018). Consequently, coordinated post-natal growth of the muscle-tendon structures in tandem with skeletal growth is critical to maintain full range of motion. When muscle-tendon growth lags skeletal growth, increased tensioning develops across the joint due to fixed anchoring of tendon to bone. This results in joint contracture with restricted range of motion. Joint contractures can occur from a variety of causes including congenital etiologies (arthrogryposis, club foot) and acquired neuromuscular causes (cerebral palsy, brachial plexus palsy, spinal muscular atrophy) that consistently arise with muscle-tendon unit (MTU) shortening(Ippolito and Ponseti, 1980; Lieber and Fridén, 2019; Mercuri et al., 2004; Nikolaou et al., 2011; Oishi et al., 2019). Among children with neuromuscular disease, development of neuromuscular contractures (NC) is common with an estimated prevalence of 17M people worldwide and an annual cost of $2.2 billion in the US(Foad et al., 2008; Verhaart et al., 2017; Yeargin-Allsopp et al., 2008). Although very little is known regarding the regulation of MTU growth by neuromuscular signaling, recent studies have elucidated changes in muscle fibrosis and inflammation following neuromuscular disease that contribute to NC formation(Graham et al., 2016; Nikolaou et al., 2011). *In vivo* biomechanical tests suggests that muscle-specific changes do not fully account for the degree of NC severity suggesting a possible role for extra-muscular tissues, such as tendon, in NC formation(Lieber and Fridén, 2019; Nikolaou et al., 2014). Despite this, little is known regarding the role of tendon in NC formation.

Tendons are dense connective tissues that transmit forces from muscle to bone to produce joint motion. During post-natal development, tendon undergoes robust proliferation and extracellular matrix deposition that constitute the strong tensile properties of tendon(Ansorge et al., 2011; Grinstein et al., 2019). Therefore, it is possible that disruption of in tendon growth may contribute to NC formation. Despite this, the role of tendon in neuromuscular contracture remains unclear. To determine whether changes in tendon elongation contribute to NC formation we established a novel surgical model of NC formation following neonatal sciatic nerve transection (SNT) in the hindlimb. Remarkably, mice developed joint NC and skeletal deformity that closely modeled human NC. Further, transcriptional changes in our model were concordant with changes in tendon from patients with NC. Using this model, we established a novel role of intramuscular myotendon elongation following neonatal SNT that constrains muscle elongation resulting in joint contracture. We further found increased myotendinous Smad4 activity in mouse and human samples of NC, with rescue of NC formation following inhibition of BMP-mediated Smad4 signaling with a small molecule inhibitor. These results suggest a novel role of Smad4 mediated myotendon elongation in NC formation.

## RESULTS

### Neonatal sciatic neurotomy: a novel model of neuromuscular contracture in the hindlimb

Hindlimb sciatic nerve transection (SNT) was performed in post-natal day 5 (P5) mice to induce neuromuscular contracture formation. Following denervation, mice were able to feed with minimal restraint. To confirm surgical denervation did not induce a local inflammatory response within the tendon, we performed flow cytometry to characterize inflammatory cell recruitment. Flow cytometry demonstrated ≤2% immune cell recruitment in denervated or sham operated tissues (Fig. S1). There was no difference in T-cell recruitment, nor differences in overall macrophage recruitment (Fig. S1). Overall, these data indicate a negligible local inflammatory response to proximal nerve transection.

**Figure S1:**
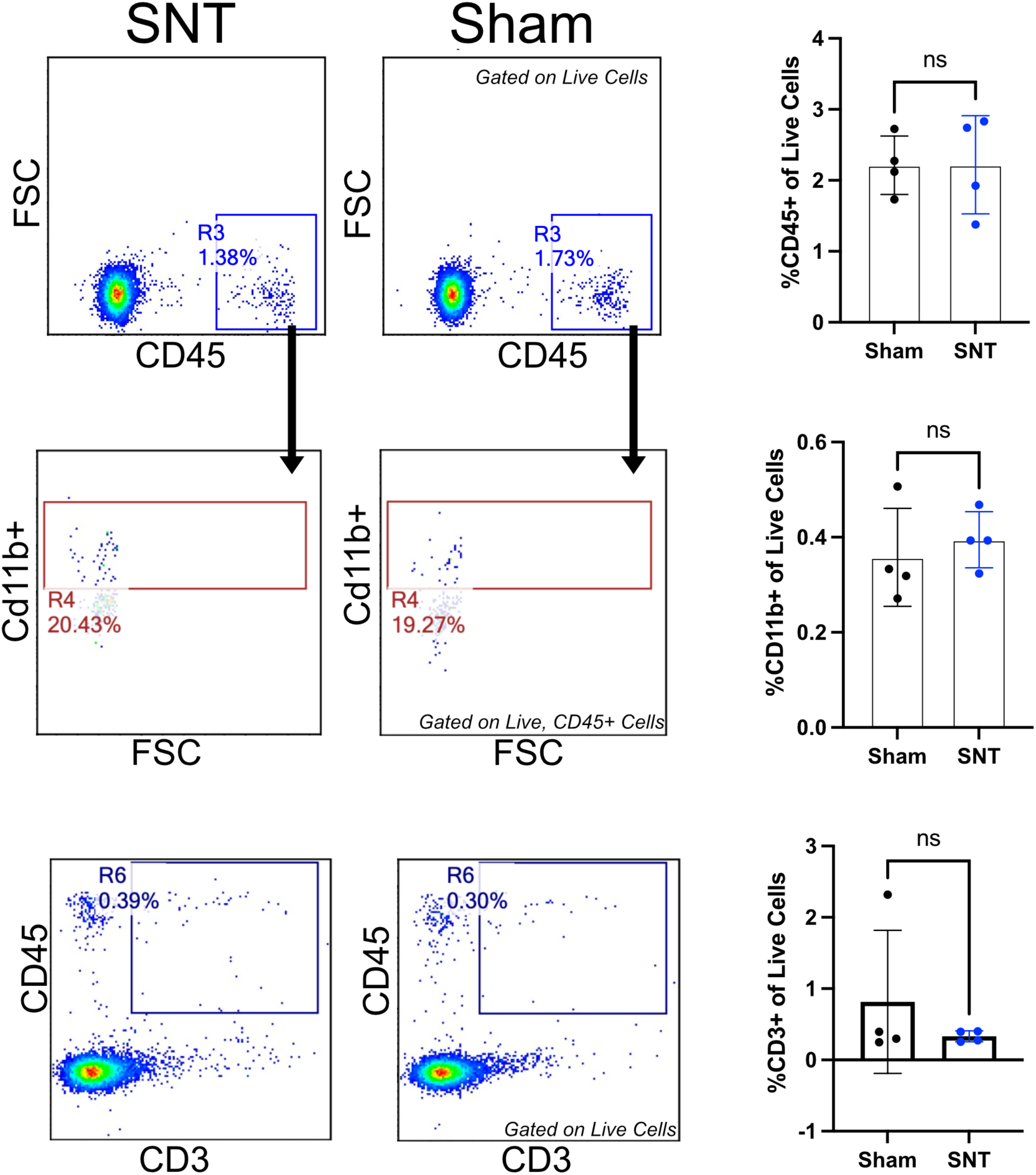
Characterization of immune cell infiltrate in tendon following denervation. Flow cytometry profiling of CD45+ immune cells in Achilles tendon 14 days following SNT or sham operation with subsequent quantification of CD45+, CD11b+ macrophages and CD3+, CD45+ T-cells. All gating was performed on singlet, DAPI-live cells (n=4 mice, 2-tailed Student’s t-test; ns, *p*>0.05).

To assess the relevance of our model to human neuromuscular contracture, we characterized the development of equinus contracture (fixed ankle plantar flexion) which commonly occurs in patients with neuromuscular disease (Greene, 2000). Under passive ankle dorsiflexion, fixed equinus contracture was noted by 14 days post-neurotomy (DPN) with widening of the tibia-midfoot angle that was sustained at 28DPN compared to sham operated controls (Fig. 1A). Restricted range of motion can occur following muscle-tendon shortening or from capsular adhesions (Kaneguchi et al., 2017; Suwankanit and Shimizu, 2022). To test this, ankle range of motion was re-assessed following myotomy which showed rescue of ankle contracture, indicating that contracture formation is primarily driven by muscle-tendon shortening and not from capsular adhesions (Fig 1B).

**Figure 1:**
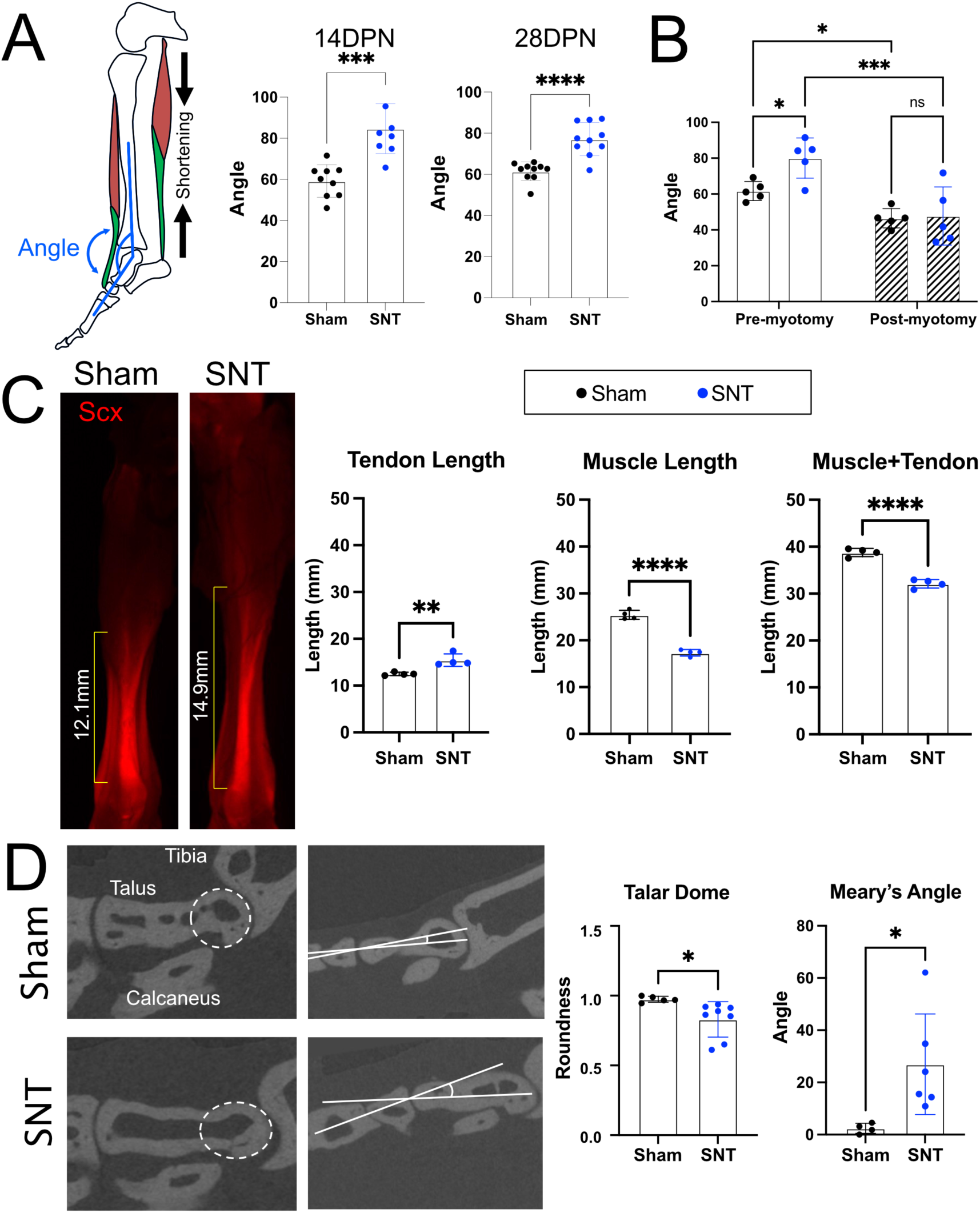
Neonatal sciatic nerve transection functionally recapitulates neuromuscular contracture formation. A) Characterization of ankle equinus was quantified by measuring the tibia-midfoot angle under passive ankle dorsiflexion at 14- and 28DPN (n=7-10 mice, 2-tailed Student’s t-test; ***, *p*<0.001). B) Assessment of ankle equinus pre- and post-triceps surae release at 28DPN (n=5 mice, 2-way ANOVA, Sidak’s post-hoc; *, *p*<0.05; ***, *p*<0.001). C) Quantification of tendon, muscle, and muscle+tendon (MTU) length in mouse hindlimbs via whole mount microscopy in Scx-mCherry transgenic mice at 14DPN (n=5 mice, 2-tailed Student’s t-test; **, *p*<0.01; ****, *p*<0.0001). D) Assessment of ankle cavus (Meary’s angle) and talar dome flattening from reconstructed micro-CT slices at P120 (115DPN) (n=4-8 mice, 2-tailed Student’s t-test; *, *p*<0.05).

We next sought to characterize changes in muscle and tendon lengthening following NC formation. Using *Scx-mCherry* reporter mice to specifically label tendon, whole mount imaging of hindlimbs at 14 days-post sciatic nerve denervation (14DPN) revealed tendon elongation with muscle shortening (Fig. 1C) (Pryce et al., 2007). Collectively, this resulted in cumulative shortening of the muscle-tendon unit. There was no difference in tibia or femur lengths following denervation.

**Figure S2:**
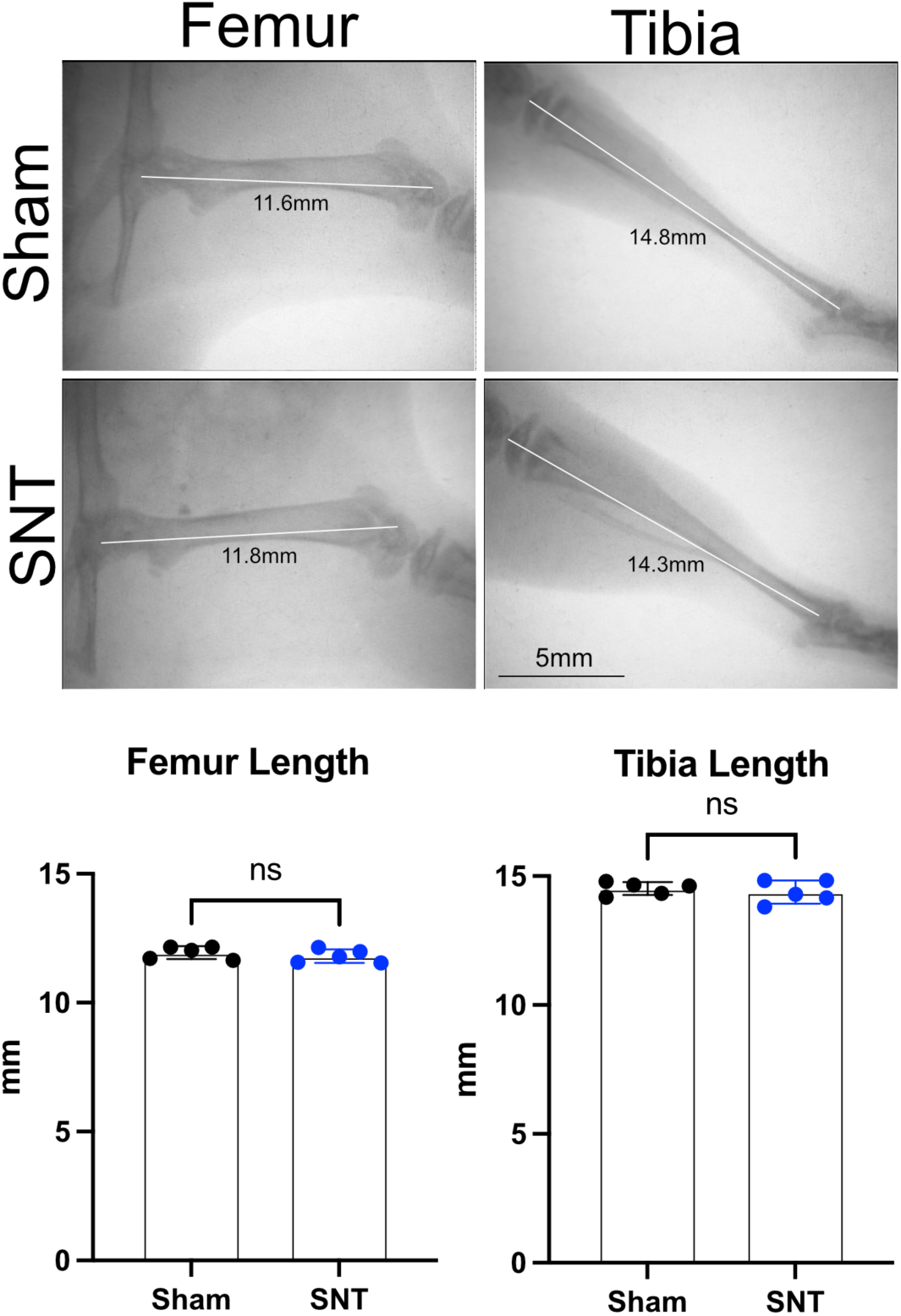
SNT does not altern hindlimb skeletal growth. X-ray radiographs of femur and tibia 28 days following sham or SNT (n=5 mice, 2-tailed Student’s t-test; ns, *p*>0.05).

Given the changes in soft-tissue growth, we next determined if long-term changes in soft-tissue tensioning may result in skeletal deformity of the foot and ankle. To quantify degree of deformity, we measured two radiographic parameters of foot and ankle dysplasia, namely degree of talar dome roundness and Meary’s angle to assess food cavus which both occur in humans with neuromuscular contracture (see Methods). Using these parameters, we analyzed *ex vivo* micro-CT reconstructions of hindlimbs in skeletally mature mice at P120 following sciatic denervation or sham operation performed at P5. Results showed development of skeletal dysplasia with talar dome flattening and increased Meary’s angle, representative of worsening cavus foot deformity (Fig 1D).

### Neonatal denervation results in altered tendon mechanical and structural properties

To characterize changes in tendon material properties we performed uniaxial mechanical testing of tendons at 28DPN following SNT or sham surgery. Tendons from denervated limbs displayed decreased tensile stiffness, maximum force, and toughness (Fig. 2A). Given the striking difference in mechanical properties we next determined changes in tendon composition. To assess changes in collagen structure, we performed picrosirius red staining of longitudinal paraffin sections along the tendon. Tendons in sham operated and sciatic denervated mice at 28DPN were uniformly collagenous with no evidence cellular infiltrates (Fig. 2B). Notably, tendons from denervated limbs were thinner with decreased collagen content from hydroxproline quantification (Fig. 2C). To characterize changes in collagen ultrastructure, we performed transmission electron microscopy (TEM) of transverse sections through the tendon midsubstance (Fig. 2D). Fibril diameters from tendons in sham operated hindlimbs were normally distributed with ∼75% of fibrils between 60-160nm in diameter. In contrast, fibrils from denervated hindlimbs had a bimodal distribution, with fewer medium sized fibrils (60-160nm, ∼59%), and greater smaller (<60nm, ∼29%) and larger (>160, ∼12%) fibrils. Despite this, there was no difference in collagen fibril packing efficiency. These results demonstrate differences in post-natal tendon maturation and growth following neonatal SNT.

**Figure 2.**
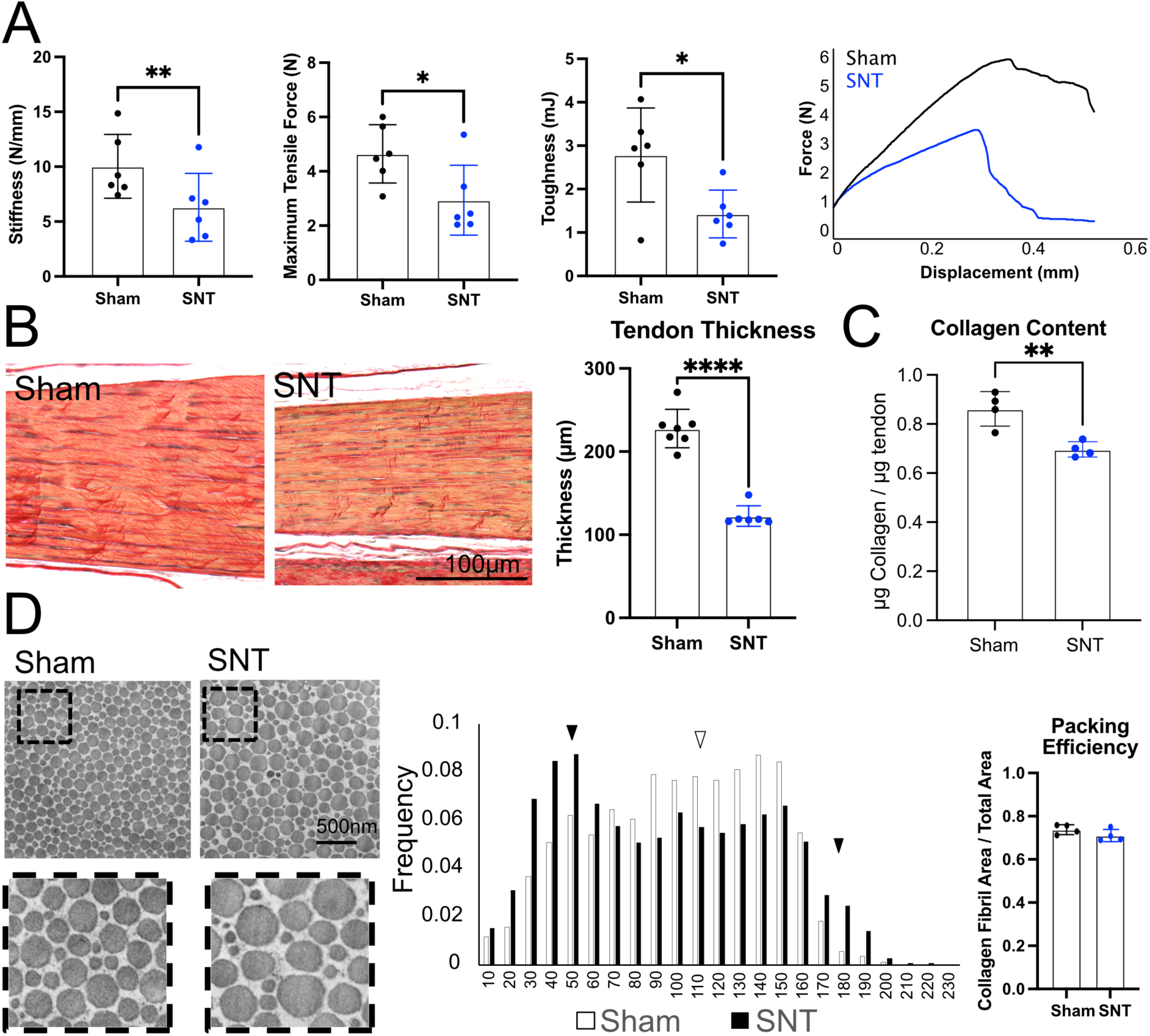
Mechanical, compositional, and structural properties of tendon following SNT. A) Characterization of tendon tensile stiffness, maximum tensile force, and toughness 28DPN (n=6 mice, 2-tailed Student’s t-test; *, *p*<0.05; **, *p*<0.01). B) Picrosirius red staining of Achilles tendon at 28DPN from longitudinal paraffin sections (n=6-7 mice, 2-tailed Student’s t-test; ****, *p*<0.0001). C) Quantification of tendon collagen composition normalized to tissue total protein in Achilles tendon from SNT or sham operated limbs collected at 28DPN (n=4 mice, 2-tailed Student’s t-test; **, *p*<0.01). D) TEM micrographs through transverse sections of the Achilles tendon midsubstance at 28DPN. Histogram demonstrating distribution of collagen fibril diameter from sham (white) and SNT (black) hindlimbs. Arrowheads represent increased frequency of fibril diameters in sham (white) or SNT (black) hindlimbs. Packing efficiency quantified as cumulative sum of collagen fibril area divided by total area (n=4 mice, 2-tailed Student’s t-test; ns, *p*>0.05).

### Tendons following SNT display a distinct neuromuscular contracture transcriptional signature

To determine transcriptional differences that may underlie structural and compositional changes in tendon following SNT, bulk RNA-seq was performed on dissected muscle and tendon tissue from SNT and sham operated limbs 14 days post-surgery. Following PCA analysis, samples from each respective group clustered closely with greater separation between sham and denervated tendon groups compared to muscle, suggesting greater transcriptomic variance among tendon groups (Fig. 3A-B). Analysis of differentially expressed genes (DEG) in tendon identified gene ontology (GO) pathways associated with collagen matrix, cell adhesion, and integrin complex regulation (Fig. 3C). In contrast, muscle GO terms were associated with endocytic and phagocytic processes (Fig. 3D).

**Figure 3:**
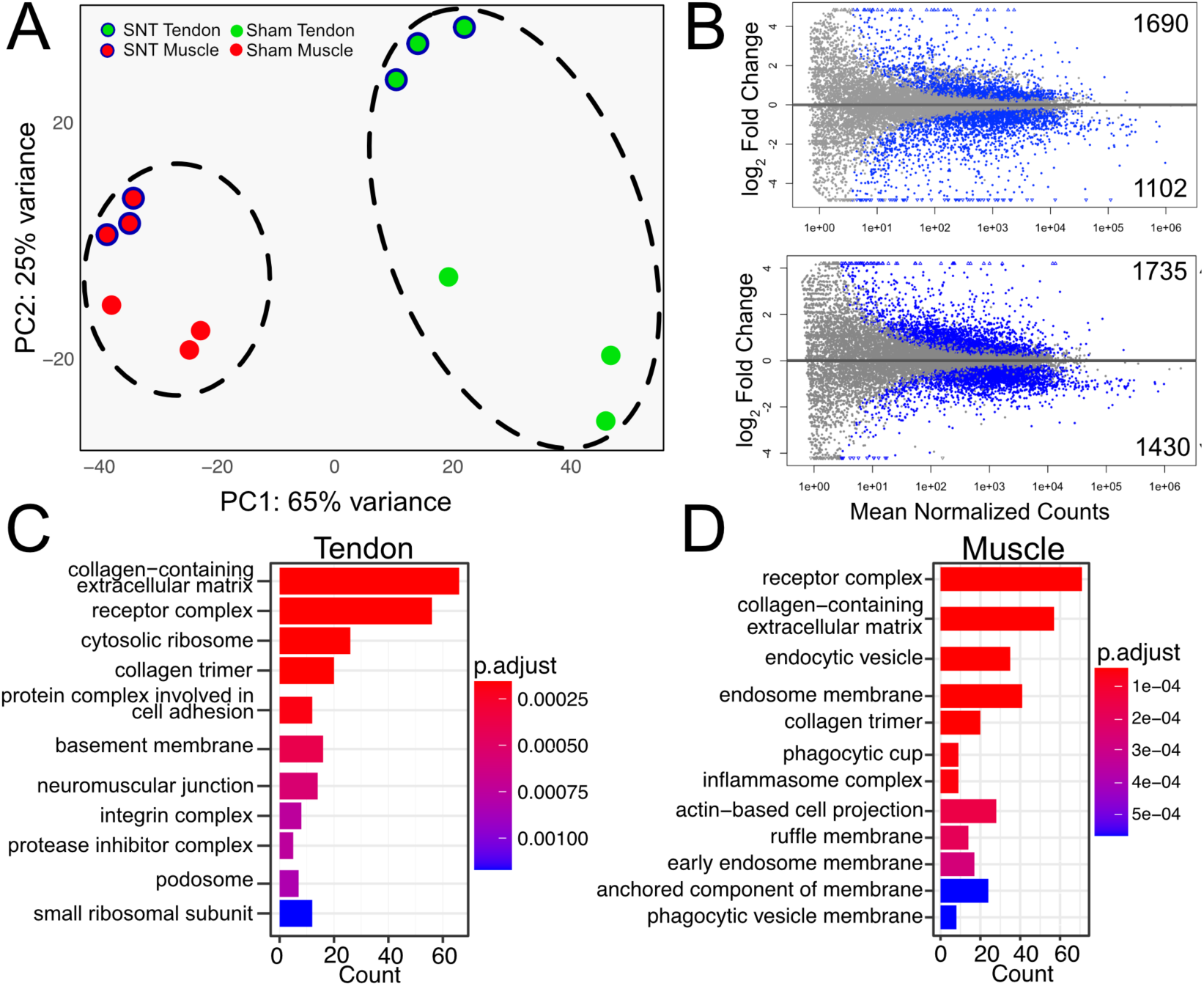
Tendon and muscle demonstrated transcriptomic changes following neonatal SNT. A) Principal component analysis (PCA) of RNA-seq transcriptomic data from mouse tendon (green) and muscle (red) in sham (no outline) operated and SNT (blue outline) hindlimbs (n=3). B) Volcano plots of normalized counts vs log_2_-fold change, with differentially expressed genes (DEG) (*p-adj* < 0.05) identified in blue marker. Counts represent number of DEGs. C) Gene ontology analysis of DEGs in tendon and in D) muscle.

To determine gene sets that may be involved in contracture formation, we next compared DEGs in mouse to DEGs from published RNA-seq of tendon tissue from human patients with tetraplegic cerebral palsy compared to typically developing children (Nemska et al., 2023) (Fig. 4A). Remarkably, gene set enrichment analysis (GSEA) of DEG in mouse tendon following SNT, revealed enrichment for genes associated with the human tetraplegic tendon gene signature (Fig. 4B). We next determined whether tendon-specific genes follow similar expression profiles in mouse and human. Real-time qPCR showed increased expression of all tendon markers (*Scx, Mkx, Tnmd*) in mouse tendon following SNT compared to sham operated limbs (Fig. 4C). In comparison, transcripts from RNA-seq of tendon markers in human tendon tissue from tetraplegic CP patients showed similar increased counts in *Scx, Mkx,* and *Tnmd* (Fig. 4C).

**Figure 4:**
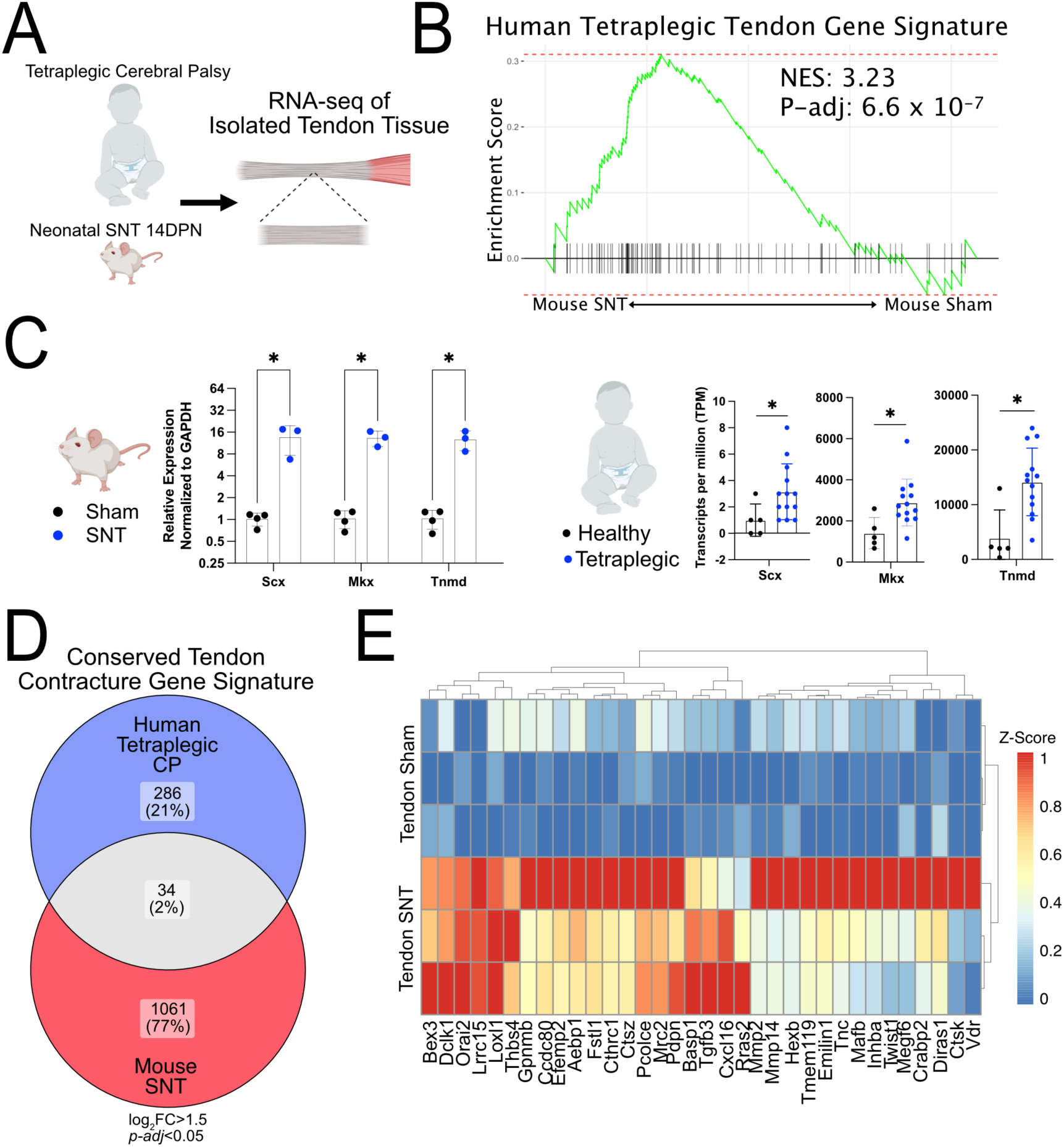
Transcriptional changes in mouse tendon following SNT recapitulate gene expression change in human tendon from patients with NC from spastic cerebral palsy. A) Schematic of experimental design. B) Gene set enrichment analysis (GSEA) for DEGs genes expressed in tendon from patients with cerebral palsy with NC is highly correlated to transcriptomic changes in mouse tendon following neonatal SNT. NES, normalized enrichment score. C) qPCR analysis of tenogenic genes in mouse tendon following sham or SNT 14DPN and transcripts per million (TPM) from published RNA-seq data of tendon tissue from typically developing children and children with NC from CP (Nemska et al., 2023) (n=6 mice, n=5-13 patients; 2-tailed Student’s t-test; *, *p*<0.05).

We next performed comparative analysis between differentially upregulated genes between mouse and human and identified 34 genes that constitute a conserved tendon contracture gene signature (Fig. 4D-E). Genes identified are associated with extracellular matrix (ECM) assembly (*Tnc, Emilin1*), ECM remodeling (*Mmp2, Mmp14, Loxl1, Pcolce*), cell adhesion (*Thbs4, Gpnmb*), and cell migration (*Pdpn, Twist1, Mrc2*) (Bressan et al., 1993; Chiquet-Ehrismann and Tucker, 2011; Del Buono, 2013; Frolova et al., 2014; Gai et al., 2014; Han et al., 2021; Herchenhan et al., 2015; Martín-Villar et al., 2006; Steiglitz et al., 2006; Yang et al., 2012).

### Myotendon elongation results in neuromuscular contracture formation

Since contracture formation occurs due to relative shortening of soft tissues with respect to bony growth, we initially hypothesized that tendon shortening might contribute to contracture formation. Despite this, we paradoxically observed tendon elongation with increased tendon compliance which would be expected to lessen degree of contracture. Further analysis of the conserved tendon contracture gene signature revealed increased expression of *Thbs4,* which is known to localize at the myotendinous junction and is required for proper muscle-tendon attachment (Subramanian and Schilling, 2014). Therefore, we hypothesized that contracture formation following SNT may result in changes at the myotendinous junction.

To test this, we first assessed extramuscular tendon (EM-tendon) and myotendon length following denervation (Fig. 5A-B). Strikingly, longitudinal picrosirius red sections of mouse hindlimbs demonstrated increased myotendon lengthening into the muscle belly in denervated limbs (Fig. 5A). Quantification of EM-tendon and myotendon lengths (Fig. 5B) following MTU dissection with muscle stripping revealed myotendon lengthening with no change in EM-tendon length (Fig. 5C). This suggests increased myotendon interposition in muscle may contribute to contracture formation by restriction of muscle elongation. To test this intramuscular myotenotomy was performed following onset of NC formation (Fig. 5D). Myotenotomy in sham operated limbs did not result in a change in ankle range of motion indicating that in typically developing mice, myotendon interposition does not significantly constrain ankle range of motion (Fig 5E). In contrast, myotenotomy in SNT limbs rescued ankle range of motion to that of sham operated limbs with or without myotenotomy. Following myotenotomy, complete Achilles tenotomy resulted in complete release of MTU restraint on ankle range of motion in both denervated and sham operated hindlimbs, with the remaining ankle motion restricted by capsular ligaments (Fig 5E). These data suggest that myotendon lengthening results in contracture formation.

**Figure 5:**
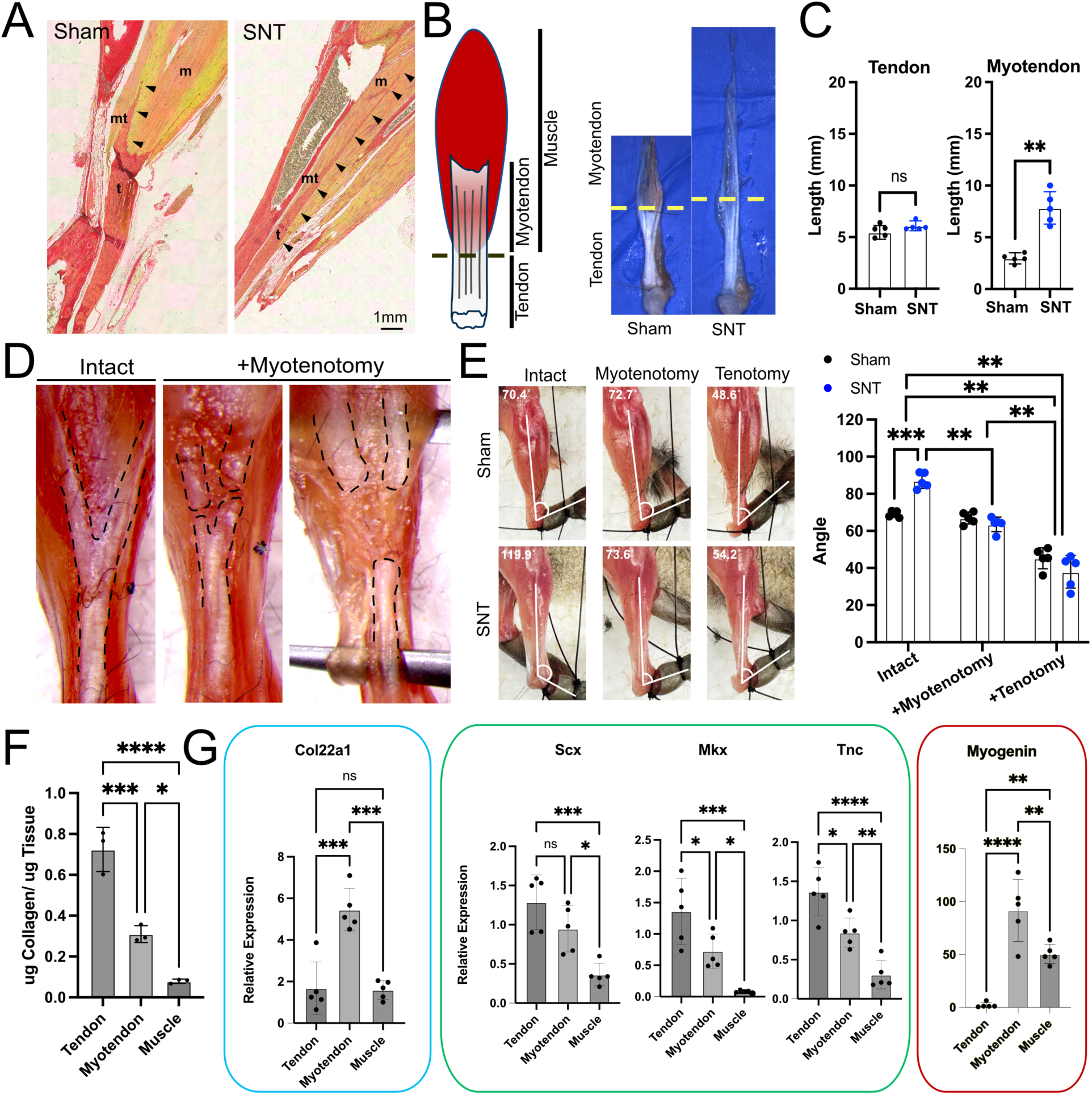
Myotendon elongation following neonatal SNT results in NC formation. A) Longitudinal paraffin sections from sham and SNT hindlimb at 28DPN stained with Picrosirius Red (t, tendon; mt, myotendon; m, muscle; black arrow heads demarcate intramuscular myotendon). B) Schematic and whole mount imaging of dissecting triceps surae following muscle stripping with dashed line differentiating tendon and myotendon transition. C) Quantification of tendon and myotendon lengths following sham operation or SNT at 28DPN (n=5 mice; 2-tailed Student’s t-test; ns, *p*>0.05, **, *p*<0.01). D) Whole mount images of mouse hindlimbs at 28DPN following SNT outlining tendon/ myotendon (left), and following intramuscular myotenotomy (middle), with intact triceps surae musculature (right). E) Photographs and bar graph demonstrating degree of ankle equinus contracture following sham operation or SNT at 28DPN before and after intramuscular tenotomy or complete Achilles tendon tenotomy. Overlays depict tibia-midfoot angle (degree of ankle equinus) in passive ankle dorsiflexion (n=5 mice, 2-way ANOVA, Sidak’s post-hoc; **, *p*<0.01; ***, *p*<0.001). F) Quantification of extra-muscular tendon (tendon), myotendon, and muscle collagen content collagen content normalized to total tissue protein from tissue following SNT collected at 28DPN (N=3-4 pooled tissues per sample; n=3 samples; 1-way ANOVA with Tukey’s post-hoc test; *, *p*<0.05; ***, *p*<0.001; ****, *p*<0.0001). G) qPCR analysis of tissue samples collected at 14DPN and assessed for markers of myotendon (blue outline), tendon (green outline), and muscle (red outline) (n=5 mice; 1-way ANOVA with Tukey’s post-hoc test; ns, *p*>0.05; *, *p*<0.05; **, *p*<0.001; ***, *p*<0.001; ****, *p*<0.0001).

To determine tissue-specific differences between EM-tendon, myotendon, and muscle from denervated limbs we performed hydroxyproline assays which demonstrated that myotendon collagen composition was in between tendon and muscle (Fig 5F). As expected, expression of the myotendinous marker *Col22a1* was restricted to the myotendon, whereas expression of tendon-specific genes (*Scx, Mkx, Tnc*) was greater than muscle but less that EM-tendon (Fig. 5G). Surprisingly, expression of the muscle marker *Myog* was elevated in the myotendon compared to muscle. These data demonstrate a unique composition and transcriptional profile of the myotendon.

### Muscle imbalance is not a principal cause of neuromuscular contracture

One hypothesis for the cause of contracture formation is due to tonic posturing resulting from asymmetric muscle loading across the joint (Chuang et al., 1998; Ho et al., 2010; Martinez-Lozano et al., 2022; Waters, 2005; Weekley et al., 2012). To test this hypothesis, we performed selective tibial (TNT) and peroneal nerve transections (PNT) to selectively denervate the triceps surae and tibialis anterior muscles, respectively (Fig. 6A). As expected, at 14DPN ankle dorsiflexion was unrestricted in TNT limbs due to muscle imbalance with unopposed tone from tibialis anterior outcompeting the atonic triceps surae (Fig. 6B). The reverse was observed in PNT limbs (Fig. 6B). If contracture formation is due to tonic posturing from muscle imbalance, then following TNT unopposed tone of the tibialis anterior would result in triceps surae lengthening with calcaneus deformity (fixed foot dorsiflexion). Similarly, the opposite would be expected in PNT limbs resulting in an equinus deformity (fixed foot plantarflexion). Paradoxically, at 28DPN ankle equinus and calcaneus deformity was observed in TNT and PNT limbs, respectively (Fig. 6B-C). At 28DPN, dissection of the Achilles tendon with muscle stripping revealed no difference in EM-tendon lengths across conditions with increased myotendon lengths following TNT and SNT (Fig. 6D-E). Taken together, these results contradict muscle imbalance as a sole cause of neuromuscular contracture formation and support myotendon lengthening following corresponding muscle denervation as a cause of neuromuscular contracture formation.

**Figure 6:**
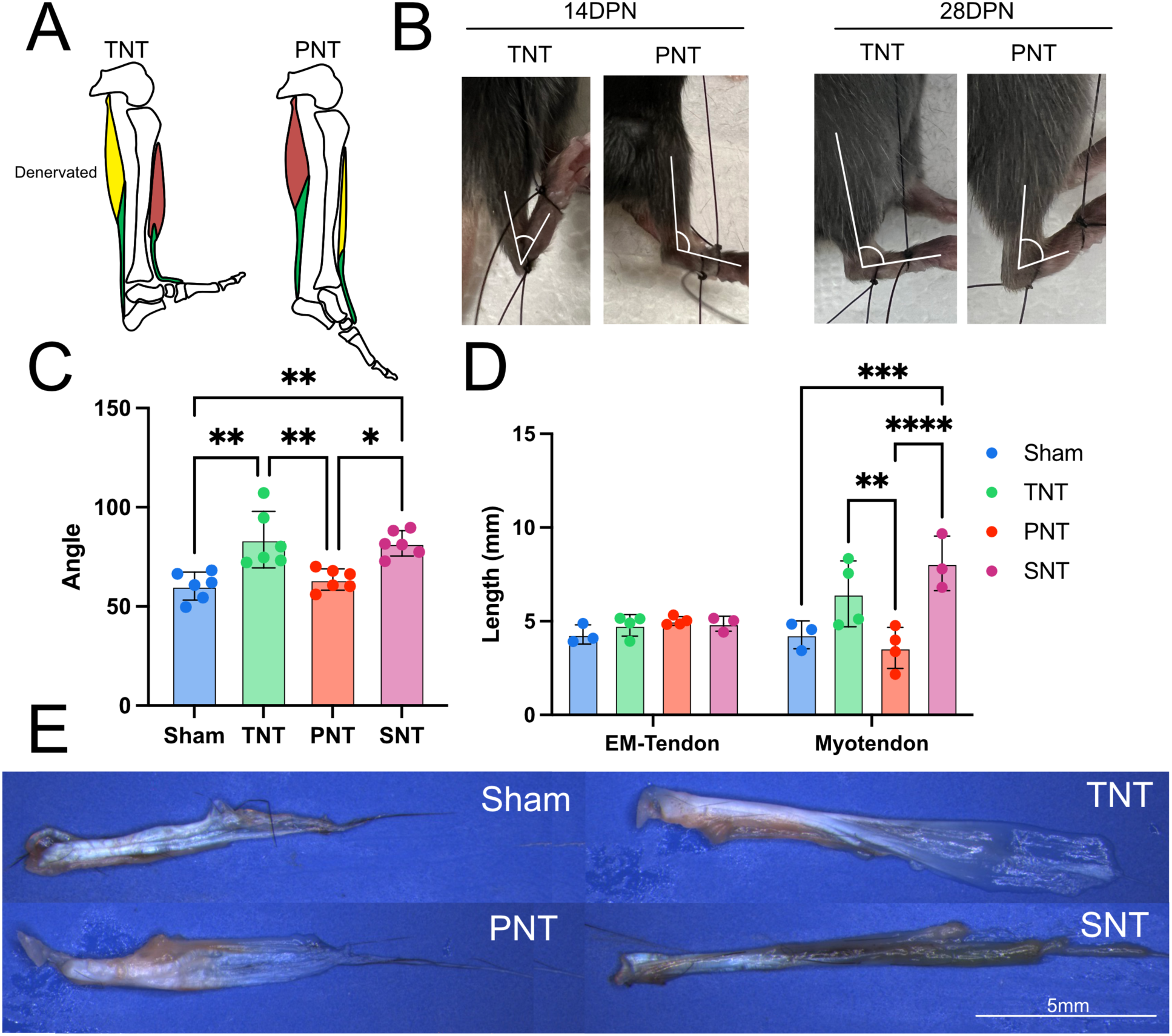
Muscle imbalance alone does not fully account for NC formation. A) Schematic demonstrating fixed ankle dorsiflexion or fixed ankle plantarflexion following selective transection of the tibial nerve or peroneal nerve, respectively (TNT, tibial nerve transection; PNT, peroneal nerve transection). Photographs demonstrating hindlimbs in passive dorsiflexion with overlays representing tibia-midfoot angle or degree of ankle equinus at 14- or 28DPN following TNT or PNT. C) Quantification of ankle equinus at 28DPN (n=6 mice; 1-way ANOVA with Tukey’s post-hoc testing; *, *p*<0.05; **, *p*<0.01). D) Quantification of EM-tendon and myotendon lengths from whole mount micrographs of dissected tissues (E) (n=3-4 mice; 2-way ANOVA with Sidak’s post-hoc testing; ** *p*<0.01; *** *p*<0.001; **** *p*<0.0001).

### Myotendon elongation following SNT occurs during a limited peri-natal window

The Achilles tendon undergoes dramatic structural changes during early post-natal growth (Ansorge et al., 2011; Grinstein et al., 2019). Therefore, to determine if aberrant myotendon elongation was a result of peri-natal denervation we performed timed SNT at P7, P14, and P12 (Fig. 7A). Mice were euthanized 28DPN. Remarkably, restricted myotendon lengthening (Fig. 7B-C) and ankle range of motion (Fig. 7D) was observed following SNT at P7 and P14, but not at P21 when compared to sham operated limbs. In contrast, there were no differences in EM-tendon lengths between SNT and sham operated limbs across all time points. These results indicate that early (<P21) neuromuscular signaling is critical for proper myotendon elongation.

**Figure 7:**
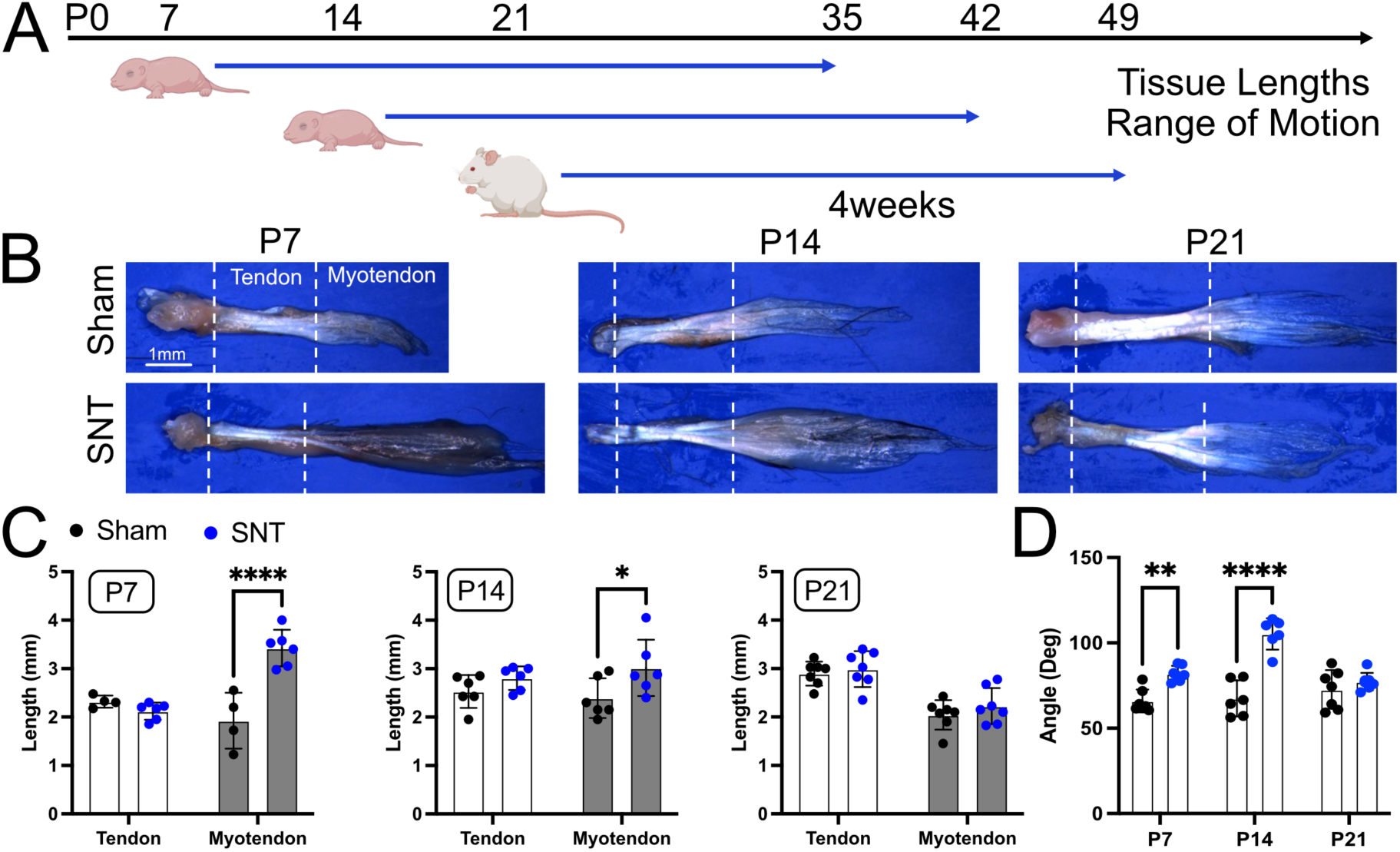
Myotendon elongation with NC formation occurs following neonatal SNT during a unique perinatal window. A) Schematic illustrating experimental design of timed SNT. B) Whole mount micrographs of dissected tissue with dashed lines indicating calcaneus insertion (left) and EM-tendon-myotendon transition (right). C) Quantification of EM-tendon and myotendon lengths and D) degree of ankle equinus contracture (n=4-6 mice; 2-way ANOVA with Sidak’s post-hoc test; *, *p*<0.05; **, *p*<0.01; ****, *p*<0.0001).

**Figure 8:**
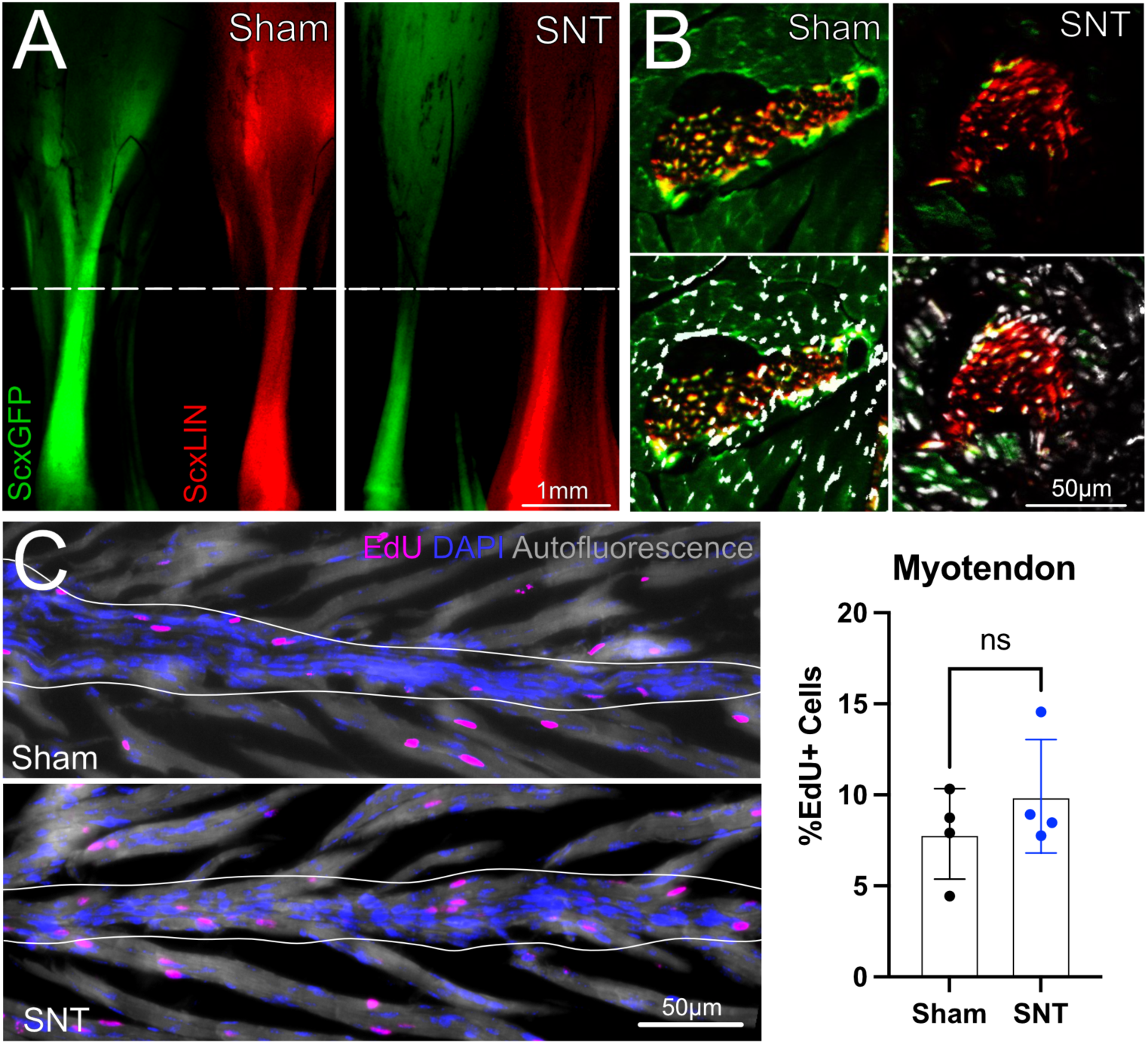
Myotendon elongation following neonatal SNT is a result intrinsically derived tenocyte infiltration. A) Whole mount micrographs of hindlimbs collected 14 days following sham operation or SNT with administration of tamoxifen at P2 in ScxGFP; ScxCreERT2; RosaT mice. Dashed line indicates EM-tendon-myotendon transition. B) Fluorescence microscopy of transverse sections at the level of the myotendon in mouse hindlimbs following sham operation or SNT 14DPN. C) EdU labeling of proliferating cells in longitudinal cryosections along the myotendon in sham and SNT operated hindlimbs collected 14DPN (n=4 mice; 2-tailed Student’s t-test; ns, *p*>0.05).

### Myotendon lengthening is driven by recruitment of intrinsic tenocytes

To determine the source of cells involved in myotendon elongation we next generated ScxGFP; ScxCreERT2; Rosa26-TdTomato (RosaT) mice to lineage trace tenocytes with tamoxifen administration at P2/ P3. We hypothesized that cells contributing to myotendon elongation originated from intrinsic tenocytes since myotendinous cells express *Scx* and are specified from migrating tenocytes during development (Schweitzer et al., 2010). Indeed, in both sham and SNT limbs, myotendon was RosaT+ indicating intrinsic tenocyte origin. In sham operated limbs, myotendon was ScxGFP+ indicating a tenocyte fate. In contrast, in SNT limbs, myotendon elongation was driven by infiltration of RosaT+, ScxGFP-cells. To determine if myotendon elongation results from increased cell proliferation we performed proliferation pulse chase experiments, with administration of EdU 3DPN at P8 during peak tendon proliferation (Grinstein et al., 2019). Analysis demonstrated no significant difference in percentage of EdU+ myotenocytes. Collectively, our results suggest that loss of neonatal innervation results in recruitment of intrinsic tenocytes to the elongating myotendon with proceeding loss of tenogenic identity (ScxGFP-, RosaT+).

### Myotendinous *Smad4* signaling is required for contracture formation

During post-natal tendon development Bmp ligands are expressed in tendon (Schlesinger et al., 2021). Notably, genetic deletion of Smad4 results in altered tendon growth with the formation of spontaneous limb contractures (Schlesinger et al., 2021). To determine whether alterations in *Bmp* signaling are observed in our model of neuromuscular contracture, GSEA analysis was performed of DEGs from mouse tendon following SNT which revealed enrichment for genes associated with “BMP response” (Fig. 9A). Realtime qPCR was performed from dissected EM-tendon, myotendon, and muscle to determine if changes in gene expression are localized to specific tissue. Interestingly, expression of *Bmpr1a*, and *Bmpr2* was increased in the myotendon compared to EM-tendon and muscle (Fig. 9B).

**Figure 9:**
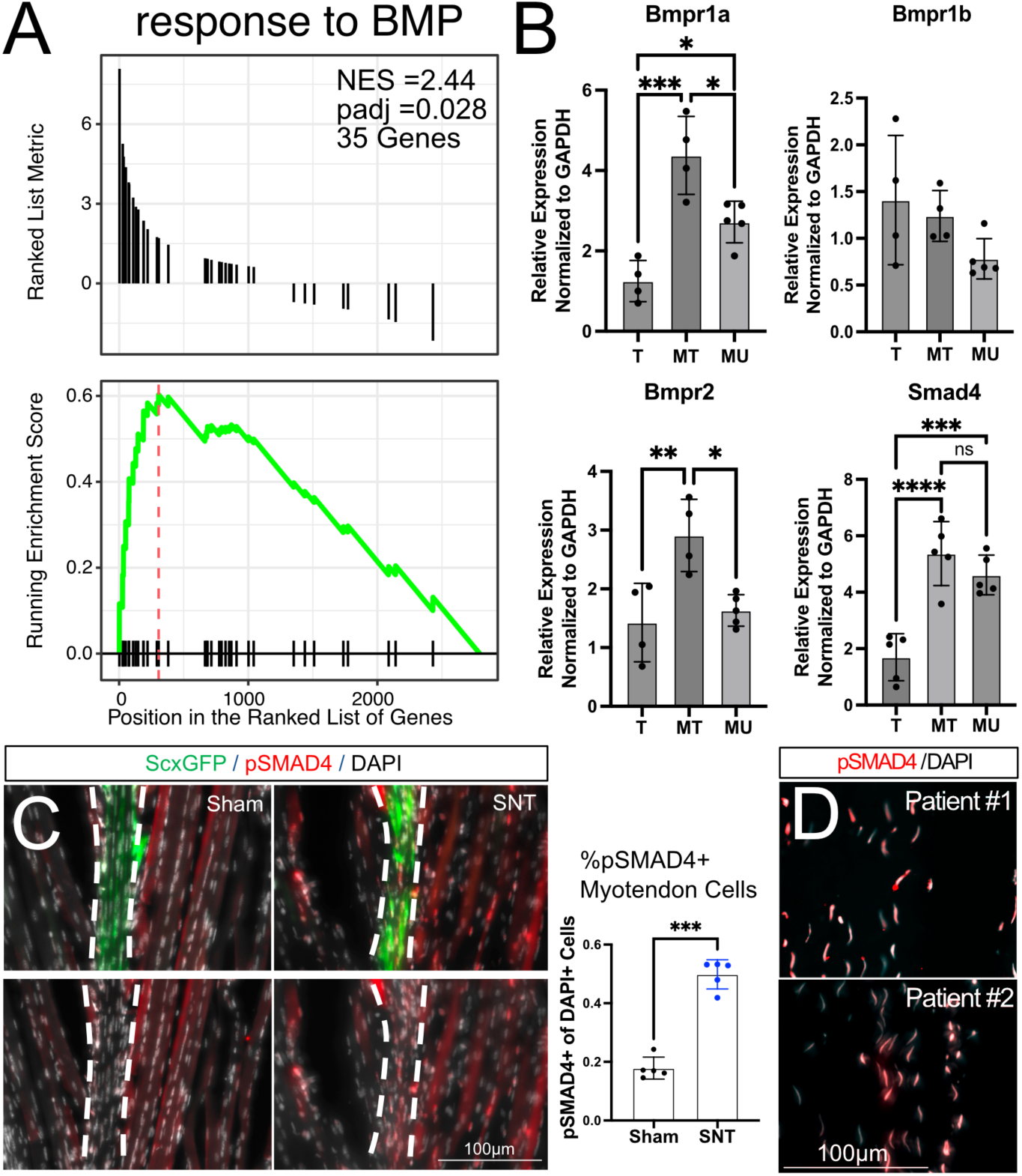
SNT results in myotendinous Smad4 activation in human and mouse NC. A) GSEA of DEGs genes expressed in tendon following SNT at 14DPN demonstrated enrichment for gene signature associated with “response to BMP”. NES, normalized enrichment score. B) Analysis of gene expression of BMP receptors and Smad4 in EM-tendon, myotendon, and muscle at 14DPN (n=4-5 mice; 1-way ANOVA with Tukey’s post-hoc test; *, *p*<0.05; **, *p*<0.01; ***, *p*<0.001; *****, *p*<0.0001). C) Immunofluorescence of longitudinal cryosections of the myotendon of mouse tissues collected at 14DPN, with quantification of pSMAD4 positive cells of total cells within the myotendon (n=5 mice; 2-tailed Student’s t-test; ***, *p*<0.001). D) Immunofluorescence of pSMAD4 from myotendon collected from human patients with cerebral palsy with NC (n=2 patients).

To determine whether Smad4 signaling is activated after SNT, we measured gene expression of Smad4 and found enhanced expression within the myotendon (Fig. 9B). Consistent with gene expression, immunofluorescence (IF) of pSMAD4 on sagittal cryosections at the MTJ demonstrated a greater proportion of myotendinous ScxGFP+ cells with nuclear SMAD4 signaling (Fig. 9C). Similarly, a greater proportion of nuclear pSMAD4 was observed in myotubules in SNT limbs compared to sham. Strikingly, IF of human tendon isolates taken at from the myotendon from patients with cerebral palsy with NC also showed evidence of nuclear pSMAD4 (Fig. 9D).

To test the requirement of *Smad4* signaling in contracture formation, we next inhibited *Smad4* signaling following. *Smad4* signaling was abrogated using the well-established small-molecule inhibitor LDN-193189 (LDN) which inhibits BMP mediated SMAD4 signaling through targeting of the ALK2, ALK3, and ALK6 receptors. LDN was administered till P21 following SNT at P5 since only SNT <P21 resulted in NC (Fig. 10A). Neonatal mice treated with LDN showed no detrimental effects and had similar growth compared to PBS treated controls (Fig. 10B). EM-tendon and myotendon lengths were decreased following LDN administration in both sham and SNT hindlimbs (Fig. 10C-D). Remarkably, LDN administration rescued contracture formation with improved ankle range of motion following SNT (Fig. 10E). Collectively, these results demonstrate expression of BMP-receptors with active Smad4 signaling in mouse and human myotendon during NC formation. Moreover, we show improvement of NC and myotendon elongation following SMAD4 inhibition in mouse.

**Figure 10:**
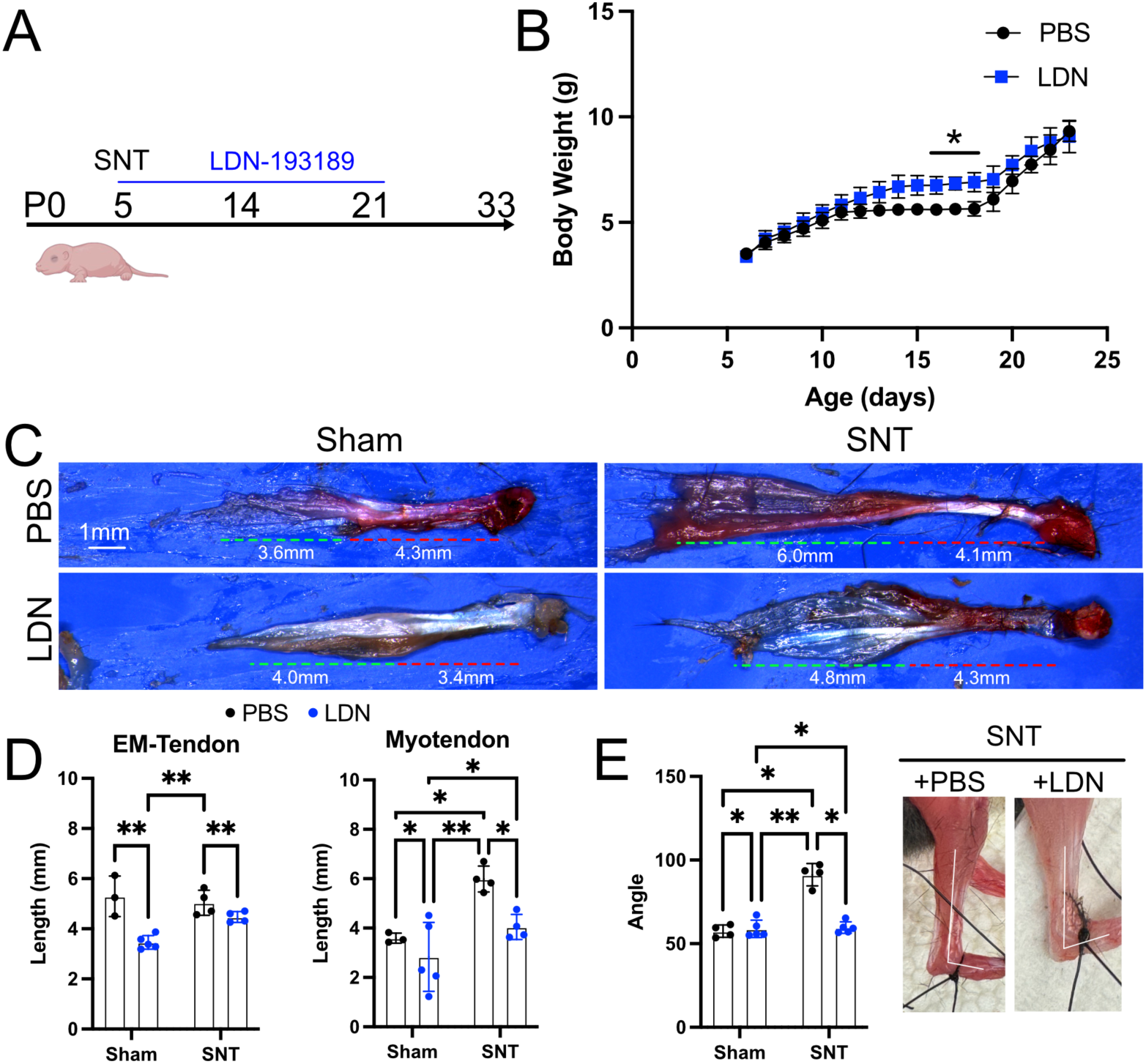
Administration of LDN resolve myotendon elongation and NC formation following neonatal SNT. A) Schematic of experimental design. B) Mouse body weight during LDN administration (n=4-5 mice; two-tailed Student’s T-test with Bonferonni correction for multiple comparisons; *, *padj*<0.05). C) Whole mount micrographs of dissected tissues collected 28DPN following sham operation of SNT with LDN or PBS administration. D) Quantification of EM-tendon and myotendon length and E) ankle equinus contracture (n=3-5 mice; 2-way ANOVA with Sidak’s post-hoc testing; *, *p*<0.05, **, *p*<0.01).

## DISCUSSION

Compromised neuronal-muscular signaling during post-natal growth results in NC from impaired MTU lengthening in relation to skeletal growth. NC occur from a variety of neuromuscular diseases including upper- (cerebral palsy) and lower- (neonatal brachial plexus injury, spinal muscular atrophy) motor neuron disease suggesting a common role of neonatal neuromuscular signaling in MTU growth (Foad et al., 2008; Verhaart et al., 2017; Yeargin-Allsopp et al., 2008). To date, several studies have observed muscle fibrosis in patients with NMD with the hypothesis that muscle fibrosis decreases muscle compliance and limits MTU lengthening with resultant joint contracture. Despite this, biomechanical studies suggest muscle fibrosis alone does not fully explain the degree of relative MTU shortening or joint contracture implicating additional causes of MTU shortening (Lieber and Fridén, 2019; Wren et al., 2010). To date, only a few studies have investigated tendon-specific changes during NC formation. Therefore, development of a murine model of NC may provide mechanistic insights into the tendon-specific contributions to NC. Further, such a model can be used to preclinically investigate the effect of novel treatments on NC progression.

We find that complete hindlimb denervation following SNT at P5 results in reproduceable and consistent ankle joint contracture formation. Similar to the clinical presentation of NC in patients with spastic cerebral palsy, we observe ankle equinus and cavus deformity, talar dome flattening, and tendon elongation with overall MTU shortening (Graham et al., 2016; Lieber and Fridén, 2019). Genetic models of spontaneous contracture formation in mouse found stochastic directionality of contracture formation with limbs either fixed in ventral or dorsal joint contracture (Schlesinger et al., 2021). In contrast, in both the clinical presentation and in our model, limbs were consistently found to be in ankle equinus (fixed plantarflexion). Collectively, these data support neonatal SNT in mouse as a clinically relevant pre-clinical model that reproduces relevant skeletal and soft-tissue deformity in NC.

Characterization of tendon mechanical, compositional, and structural properties following neonatal SNT demonstrated decreased tendon mechanical stiffness, thickness, and collagen content with an altered distribution of collagen fibril diameter. During post-natal development, collagen fibril diameter small to large fibers(Ansorge et al., 2011). Thus, is it possible that SNT results in partial arrest of fibril reorganization resulting in a bimodal distribution of fiber diameters. How disruption of neuromuscular signaling affects post-natal tendon development remains to be investigated. During post-natal development, tendon undergoes dramatic changes in material stiffness and properties (Ansorge et al., 2011). Using *in vitro* tissue engineering approaches, numerous studies have demonstrated the importance of cellular loading on tenocyte collagen deposition and organization. Therefore, one possibility is neonatal SNT may result in tendon-specific changes due to decreased muscle contractile loading. Careful comparison of load-bearing and non-load bearing tendons has identified a unique association between the onset of neonatal locomotion with the increase in material properties of load-bearing tendons (Theodossiou et al., 2019). These data support the role of contractile muscle loading in regulating post-natal development of load bearing tendons. Alternatively, it is possible that loss of neuronal signaling may independently regulate tendon maturation. Recently, it was observed that constitutive activation of *Piezo2* in proprioceptive sensory neurons that innervate muscle spindles and Golgi tendon organs resulted in shortening of tendon length with joint contracture formation (Ma et al., 2023). Investigation into whether tendon-specific changes following neonatal SNT result from disruption of neural signaling, or through loss of muscle contractile loading may help to shed light into the mechanisms that regulate post-natal tendon growth and NC formation. Moreover, distinguishing the role of afferent and efferent neuronal signaling in NC formation may further elucidate the regulation of NC formation.

Analysis of transcriptomic changes in muscle and tendon following neonatal SNT revealed dramatic changes in both tissues. Consistent with observed tendon compositional changes following neonatal SNT, gene ontology analysis revealed altered expression of pathways associated with collagen matrix regulation. In muscle, analysis revealed pathways associated with endocytosis and phagocytosis. We suspect these processes are largely activated following muscle denervation mediated catabolism (Libelius et al., 1978). In cases of muscle disuse or denervation, ubiquitin-mediated proteasomal degradation leads to atrophy and is in part attenuated by macrophage recruitment (Dumont and Frenette, 2010; Kawanishi et al., 2018; Khalil, 2018).

Gene set enrichment analysis of DEGs from tendon following neonatal SNT showed significant enrichment with tendon samples in patients with NC from tetraplegic cerebral palsy. Interestingly, these results suggest similar transcriptional changes between divergent causes of NC formation. In cerebral palsy, it is thought that disruption of upper motor neuron signaling leads to tonic spasticity leading to NC formation. In contrast, neonatal SNT results in a lower motor neuron disruption with decreased tone. In both clinical scenarios, joint contractures occur. Moreover, analysis of muscle in spastic cerebral palsy and following neonatal denervation show similar architectural changes (Nikolaou et al., 2022). Collectively, these data propose that NC from upper and lower motor neuron disease results from a common disruption in neuromuscular signaling.

Analysis of tendon structural and mechanical properties demonstrated increased tendon elongation and compliance. Paradoxically, rather than contributing to decreased MTU elongation these results suggest changes in tendon may serve to mitigate joint contracture. Further analysis of conserved DEGs genes between human and mouse tendon in NC revealed increased expression of *Thbs4*, which is known to preferentially localize to the myotendinous junction (Subramanian and Schilling, 2014). Histology and tissue dissection revealed elongation of intramuscular myotendinous tissue within the muscle belly, while compositional analysis found that myotendinous tissue had increased collagen content in comparison to muscle. Based on these observations, we propose that elongation of myotendon within muscle belly constrains muscle elongation. Interposition of stiffer myotendinous tissue within more compliant muscle, creates a composite tissue with overall stiffness predicted to be greater than that of muscle and would result in restricted muscle elongation (Bar-On and Wagner, 2013). This would also reconcile the observation that the majority of MTU shortening is resultant from muscle shortening. Consistent with this hypothesis, intramuscular myotenotomy rescued NC following neonatal SNT, permitting muscle elongation. Interestingly, intramuscular myotenotomy similarly improves joint motion in patients with NC (Altuntas et al., 2011; Dagge et al., 2012; Majestro et al., 1971).

Gene expression of EM-tendon, myotendon, and muscle confirmed specific expression of the myotendinous marker *Col22a1* within the myotendon, and expression of tenogenic genes (*Scx, Mkx, Tnc*) restricted to tendon and myotendon (Huang, 2017; Malbouyres et al., 2022). Surprisingly, expression of the myogenic transcription factor *Myogenin* was increased in myotendon. Several sources of bipotent fibromyogenic cells have been previously reported that contribute both to tendon and muscle following injury (Esteves de Lima et al., 2021; Nassari et al., 2017; Wilder Scott et al., 2019). It is possible that a bipotent source of tenomyogenic cells undergoes rapid expansion and infiltration into muscle following neonatal SNT, with incomplete fate change leading to an intermediary cell state. Whether myotendon elongation results from expansion of bipotent fibromuscular progenitors is an open question for further investigation.

One long-standing hypothesis in the neuromuscular field is that NC formation is a result of muscle imbalance, with asymmetric tone resulting in tonic posturing that results in MTU shortening due to absence of tissue loading/ lengthening (Soldado et al., 2014; Waters, 2005). For example, it has been widely observed that in children following biceps denervation due to partial neonatal brachial plexus injury, progressive paradoxical flexion contracture is observed in adolescence following initial elbow extension posturing during infancy (Kay, 1998). To address this question, using our model of neonatal nerve transection, we performed partial tibial (TNT) and peroneal (PNT) nerve transections to induce lower extremity muscle imbalance. With TNT, loss of triceps surae tone would produce ankle dorsiflexion, with the opposite effect following PNT. As expected, at 14DPN ankle joints were in apparent dorsiflexion following TNT, and plantarflexion following PNT. Surprisingly, by 28DPN reversal of joint positioning was observed with increased fixed plantarflexion of ankles following TNT and the opposite in PNT. With the tibiotalar joint as a fulcrum, this would be explained by shortening of the corresponding MTU with denervation. Analysis of triceps surae tissue showed myotendon elongation following SNT and TNT, but not with sham or PNT. These results support myotendon elongation following denervation versus tonic posturing as a cause of NC. We propose that paradoxical contracture reversal in scenarios of neonatal muscle imbalance occurs due to progressive shortening of the denervated MTU due to myotendon elongation with constraint of muscle elongation.

Previous analysis of mouse Achilles tendon found peak cellularity and turnover at <P21 with limited cell division occurring after P21 (Grinstein et al., 2019). Tendon mechanical stiffness increases dramatically after P21 compared to tendons from ≤P14, suggesting dramatic changes in collagen maturation between P14-P21 (Ansorge et al., 2011). These data suggest a unique early post-natal window during which changes in tendon growth and organization are specified. To determine whether tendon specific changes during NC are a result of disrupted early growth or persistent loss of contractile muscle loading, sequential SNT of neonatal mice at P7, P14, and P21 was performed. SNT at P7 and P14 resulted in EM-tendon elongation with joint contracture formation, while SNT at P21 did not. Notably, EM-tendon length was unchanged compared to sham operated limbs when SNT was performed at all timepoints. Consistent with our findings, deletion of Piezo2 in proprioceptive neurons resulted in joint contracture formation when performed prior to P10, but not when performed after P21 (Ma et al., 2023). Interestingly, in cases of NC in patients with spinal muscular atrophy, patients with the juvenile onset form do not develop contractures, while patients with neonatal onset do (Mercuri et al., 2020). These findings suggest that specification of myotendon-muscle interposition is determined during early (<P21) post-natal development during peak tendon growth (Grinstein et al., 2019).

Lineage tracing experiments found that myotendinous cells that infiltrate into muscle originate from intrinsic tenocytes (ScxLIN+) and are not recruited from peripheral (Scx-) sources. It is also possible that myotendinous cells arise from Col22a1+, Scx+ myotendinous cells, distinct from Col22a1-, Scx+ EM-tenocytes; however, myotendinous Cre drivers do not currently exist to tease apart this difference. Surprisingly, infiltrating myotendinous cells in SNT hindlimbs were ScxGFP-, while myotendinous tissue in sham operated limbs were ScxGFP+ indicating loss of myotendinous tenocyte identify following SNT. Notably, deletion of Tgfb2r with ScxCre similarly resulted in tenocyte dedifferentiation with loss of Scx expression (Tan et al., 2020). However, why this is restricted to myotendinous cells and not EM-tendon following SNT remains unclear. Myotendon elongation is either due to increased proliferation of myotenocytes or from migration of intrinsic tenocytes. To determine this, we performed EdU-pulse chase experiments which demonstrated no difference in myotendon proliferation between sham and SNT hindlimbs. These data support myotendon elongation is the result of migration of intrinsically derived tenocytes, not due to proliferation. Interestingly, spontaneous contracture formation following deletion of Smad4 signaling in tendon with ScxCre mice, similarly found no differences in tendon proliferation despite decreased cellularity (Schlesinger et al., 2021). While the authors hypothesized that the resultant decreased tendon cellularity may be due to decreased recruitment of tenocytes, an alternative hypothesis is that increased emigration of intrinsically derived tenocytes into the muscle, without compensatory increased proliferation may result in myotendon elongation with decreased EM-tendon cellularity. Supporting this hypothesis, we identified genes associated with migration (*Pdpn, Twist1, Mrc2*) and cell adhesion (*Thbs4, Gpnmb*) in our conserved contracture gene signature. We propose that denervation results in myotenocyte dedifferentiation with unrestricted migration into muscle resulting in contracture formation.

Although post-natal tendon elongation was found to be dependent on BMP mediated SMAD4 signaling, prior studies using the genetic *ScxCre* driver abrogated signaling in all Scx-derived cells (EM-tendon + myotendon) (Schlesinger et al., 2021). Our studies also suggest changes in response to BMP signaling during NC formation with specific upregulation of *Bmp* receptors in myotendon. In denervated muscle, release of BMP ligands autonomously limits atrophic loss of muscle mass through SMAD4 activation (Sartori et al., 2013). Therefore, as expected we observed increased pSMAD4 activity in myotubes following neonatal SNT. Surprisingly, we also observed increased pSMAD4 activity in mouse myotendon following SNT and in human myotendon from patients with NC due to spastic cerebral palsy. Since genetic deletion of *Smad4* using *ScxCre* led to global tendon shortening, we propose that increased pSMAD4 activity following neonatal SNT would lead to myotendon elongation. In confirmation of our hypothesis, administration of BMP-mediated SMAD4 inhibitor LDN, lead to reduced myotendon and tendon elongation compared to PBS treated controls. Remarkably, LDN administration also improved joint contracture. It is unclear whether direct inhibition of myotendon signaling or whether indirect inhibition through off target effects in muscle may have rescued myotendon lengthening through muscle-myotendon crosstalk. Despite this, these results suggest tendon directed therapies may offer novel therapeutic options to treat NC formation.

Here, we develop a novel surgical model of NC in mouse that demonstrates functional and transcriptional relevance to human NC. Further, we show that NC formation results, in part, due to myotendon elongation from infiltration of intrinsically derived tenocytes that lose tendon cell identity. Increased myotendon interposition results in constrained muscle elongation and contracture formation due to increased Smad4 signaling. Lastly, we find that inhibition of BMP-mediated Smad4 signaling rescues myotendon elongation and NC formation following neonatal SNT in our preclinical model of NC.

## MATERIALS AND METHODS

### Mice

ScxGP and Scx-mCherry mice were used to identify label tendon *in vivo* as previously described (Pryce et al., 2007). Scx^CreERT2^; R26^LSL-tdTomato^ (ScxCET) mice were generated for lineage tracing studies. Recombination was carried out with gavage of 25µL of tamoxifen at 10/mL in corn oil at P2 and P3 prior to nerve transection at P5. All animal studies were conducted in accordance with Institutional Animal Care and Use Committee (IACUC) at Columbia University, and all mice were housed in sterile barrier facilities operated by the Institute of Comparative Medicine at Columbia University. Equal number of male and female mice were allocated to studies by random allocation. All sample treatments and conditions were anonymized during analyses and performed in a blinded fashion.

### Nerve Transection

Neonatal mice were anesthetized by isoflurane inhalation. Mice were placed in the lateral decubitus position, and an incision was made over the femur. An avascular place was developed by splitting of the iliotibial band overlying the proximal femur. The sciatic nerve was identified and transected. To prevent peripheral nerve healing, the proximal nerve stump was resected and ∼1-2mm of nerve was excised to prevent re-apposition of nerve ends. For partial tibial and peroneal nerve transections an incision was made over the distal femur. Following splitting of the iliotibial band, the sciatic nerve was identified proximally with distal tibial, sural, and peroneal nerve continuations. Using microdissection scissors, the nerves were separated and the respective tibial or peroneal nerves were cut. Importantly, only the nerve to be cut was manipulated with microforces to prevent incidental trauma to the nerves left intact. Following transection, incision was closed with VetBond skin adhesive and animals were returned to full cage activity. Sham injuries were performed on contralateral limbs where the iliotibial band was split and nerve identified but not transected.

### FACS isolation and analysis of macrophages and Tregs

#### Tendon single cell suspension preparation

Tendon digestion and single-cell suspension preparation was performed prior to FACS isolation and analysis. At the pre-specified endpoint, tendons were dissected out and incubated in collagenase solution for 4 hours at 37C on a rocker. Collagenase solution was prepared with 5mg/mL Collagenase I (Cat. # LS004196, Worthington Biochemical) and 1mg/mL Collagenase IV (Cat. # LS004188, Worthington Biochemical) in serum free media. Following tendon digestion, tendons were triturated, spin washed (500 x *g* for 5 min at 4C) and resuspended in FACs buffer (2% FBS in PBS supplemented with in 2mM EDTA). Resuspended cells were passed through a 70μm sterile filter to force cell clumps into a single cell suspension. Single cell suspensions were then kept on ice for flow cytometry.

#### Macrophage staining

Cells in suspension were stained against CD45 (Biolegend, Cat: 10311, Pe/Cy7, 1:100) and CD11b (Biolegend, Cat: 101222, AF700, 1:100) for 30mins at 4C in the dark. Gating on CD45+ and CD11b+ cells was to select for macrophages. DAPI staining was performed for live/ dead cell identification, and DAPI+ cells were excluded as dead.

#### T-Cell Staining

Cells in suspension were stained against CD3 (ThermoFisher, Cat: 46-0032-82, PerCP, 1:100) and CD45 (Biolegend, Cat: 10311, Pe/Cy7, 1:100).

Macrophage and T-cell gates were pre-determined from cytometry of splenic cells. Following, gates established using the spleen, were applied to cells harvested from tendon. Flow cytometry was performed using the NovoCyte Quanteon (Agilent) flow cytometer at the Columbia Stem Cell Initiative Flow Cytometry Core. Flow cytometry analysis was performed using NovoExpress Software.

### RNA isolation, reverse transcription, and qRT-PCR

For RNA isolation bulk tendon, myotendon, and muscle were dissected out. Two to three specimens were combined per sample to ensure adequate RNA following isolation. Specimens were then mechanical digested by tissue homogenizer (Omni International, NY) in 1mL of Trizol reagent. Following homogenization, RNA isolation was carried out using Trizol/ chloroform extraction. After Trizol/ chloroform RNA isolation, all RNAs were quantified using NanoDrop2000. Reverse transcription was performed using SuperScript VILO (ThermoFisher, Cat: 11754050) and qRT-PCR performed using SYBR Green PCR Master Mix (ThermoFisher, Cat: 4309155). Mouse primer sequences listed below. RNA samples were collected from 3-5 independent mice and ran in triplicate.

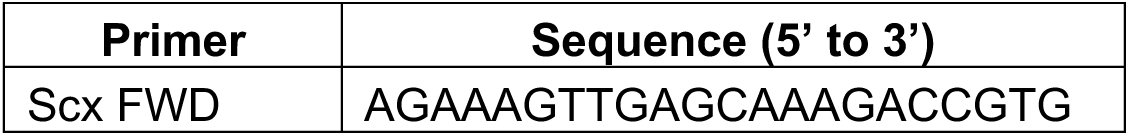
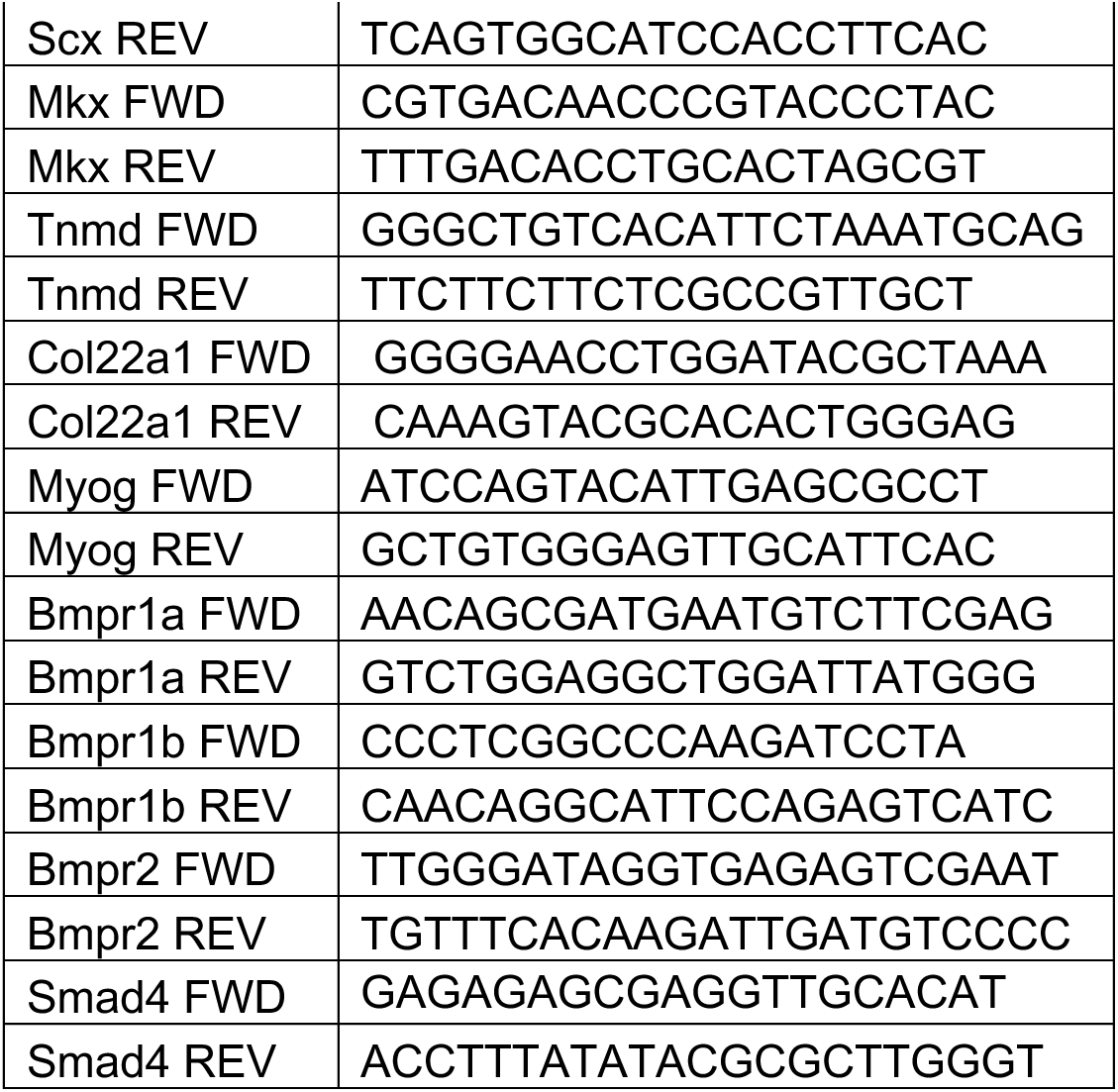

### Bulk RNA sequencing and analysis

RNA was isolated using Trizol/chloroform extraction from mouse muscle and whole tendon 14 days following either sham operation or sciatic nerve transection. RNA concentrations were measured with a NanoDrop spectrophotometer (Thermofisher), and quality was assessed with an Agilent TapeStation with DV200 ≥ 75% for all samples. RNA amplification, library preparation, and sequencing were performed by Azenta/Genewiz. Samples were sequenced on the Illumina HiSeq using a 2×150 bp sequencing. Sequence reads were trimmed to remove possible adapter sequences and nucleotides with poor quality using Trimmomatic v.0.36. The trimmed reads were mapped to the Mus musculus GRCm38 reference genome (ENSEMBLE) using HISAT2 aligner on Galaxy (PMCID: PMC9252830). Unique gene counts were calculated using featureCounts from Subread package v.1.5.2 (PMID: 24227677). Analysis was performed on R version 4.2.1. After quantification of gene counts, differential gene expression analysis was performed using DESeq2 v1.38.3. Differentially expressed genes (DEGs) were identified with Benjamini-Hochberg correction for multiple comparisons with a significance of adjusted p-value (p-adj) < 0.05 and log2 fold-change > 1.5. Bulk RNA-seq data from human tendon tissue from patients with tetraplegic cerebral palsy with ND and typically developing children was obtained from previously published data available on NCI BioProject Database (accession: PRJNA1004310) (Nemska et al., 2023).

Principal component analysis and hierarchical clustering were performed on all DEGs. Gene ontology and gene set enrichment analysis was performed on DEGs using g:Profiler (Raudvere et al., 2019). Conserved contracture gene signature was defined by first identifying mouse and human differentially upregulated genes with *padj* < 0.05 and log2 fold-change > 1.5 compared to control tendon (e.g., sham operated in mouse and typically developing in human). Intersectionality analysis was then performed to identify conserved genes that comprise a distinct and conserved contracture gene signature.

### Histology and immunofluorescence

#### Cryosectioning and Immunostaining

For immunofluorescence histology, samples were fixed in 4% paraformaldehyde overnight at 4C and then were decalcified in 0.5M EDT, replaced every 3 days, until bones were pliable. Limbs were embedded in OCT and frozen and stored at - 80C until use. Alternating transverse or longitudinal cryosections were collected at 10μm thickness, along the length of the tendon. Briefly, cells were permeabilized for 10 min with 0.1% triton X-100 in PBS followed by three PBX washes for 5 mins each. Following, slides were incubated with 10% donkey serum in 0.1% Triton X-100 in PBS for 30 min at room temperature. Immunostaining against pSMAD4 (ThermoFisher Scientific, Catalog # PA5-64712) was performed at 1:100 diluted in 1% BSA in 0.1% triton X-100 in PBS in a humidified chamber overnight at 4°C. Following three PBS washes, secondary antibody staining was performed with Cy5 (Jackson ImmunoResearch), and slides were counterstained with DAPI to visualize nuclei. For quantification, immunofluorescence microscopy (Zeiss Apotome) was performed on serial sections. Quantifications were performed across serial sections using ImageJ, and then averaged to achieve an individual result per tendon. EdU uptake and staining assays were performed as per manufacturer’s instructions (Click-iT EdU, Life Technologies).

#### Longitudinal Paraffin Sectioning and Picosirius Red Staining

For picrosirius red staining, limbs were fixed in 4% paraformaldehyde overnight at 4°C. Following fixation, limbs were decalcified in EDTA. Limbs were then dehydrated, and embedded in 75% ethanol. 6μm longitudinal sections were collected and stained with picrosirius red. Specimen sectioning and staining was performed by the Molecular Pathology and Histology Core Facility at Columbia University.

### Whole mount fluorescence imaging

Limbs were first fixed in 4% paraformaldehyde and incubated at 4°C overnight. For whole mount fluorescence imaging, skin was dissected away and tendons were visualized using a Leica M165FC stereomicroscope with filters for fluorescence.

### Micro-CT and ankle deformity assessment

For ex vivo micro-CT imaging, specimens were fixed overnight in 4% PFA in PBS. Scans were performed with a vivaCT 80 microCT scanner (Scanco). The source was set at 55KVp and 145µA. During scanning, 1000 projections over 180° (3072 samples) were recorded with a final voxel size of 10.4 µm. Slices were transformed using ImageJ (NIH) to identify a plane that was collinear with the talus and first meta-tarsal. For consistency, a sagittal slice at the midpoint of the talus was used. A line was drawn along the longitudinal axis of the talus. A secondary line was drawn through the longitudinal axis of the first metatarsal and navicular. The talus-first metatarsal angle (Meary’s angle) between these lines was measured and is a well validated measure of ankle cavus which commonly occurs in NC of the foot (Eilert, 1984). Following, degree of talar dome flattening was measured as an additional measure of skeletal deformity that is observed in long standing NC of the ankle (Dunn and Samuelson, 1974; Machida et al., 2017). Through image reconstruction in ImageJ, a sagittal slice was obtained through the midpoint and along the longitudinal axis of the talus. A circle was drawn, whose circumference aligned over the talar dome arc. Roundness of the circle was measured using the following inbuilt ImageJ formula where, roundness = 4*Area/(π*Major Axis^2^) with a value of 1 indicating a perfect circle.

### Tendon and myotendon length quantification

At the prespecified time-point, mice were euthanized and gastrocnemius muscles were released from their origin and Achilles tendon along with calcaneus insertion released from the calcaneus. Under dissection microscope, the Achilles tendon was marked at the most proximal point where muscle was noticeable (i.e. the most distal point where the first myofiber attachment to tendon is observed). Using the blunt end of a scalpel the muscle was gently scraped from the tendon until only the remaining fibrous myotendinous tissue remained. Specimen were placed in PBS until prior to length measurement. Specimen were then imaged under a Leica M165FC stereomicroscope. The distance from the calcaneus insertion to the marking represented EM-tendon length, and distance from the marking to the most proximal point represented myotendon length. Measurements were calibrated during each imaging sequence by photographing a caliper set to 5mm to convert pixel distances to mm. *Whole mount microscopy, dissected tissues*

### Ankle contracture assessment

Assessment was performed under anesthesia or immediately following euthanasia to measure ankle contracture without presence of active tone. Two slips knots were created using 4-0 nylon suture. With the mouse positioned in lateral decubitus, one slip knot was tightened over the hindfoot with care not to overlie the Achilles tendon insertion. The other slip knot was tightened over the midfoot approximately 5mm apart. A 5g weight was then hung to create a static dorsiflexion force. With the knees flexed at 90degrees, a lateral image of the ankle was taken and the angle between the tibia and midfoot was measured to quantify degree of ankle equinus contracture (fixed plantarflexion), with a worsening equinus contracture resulting in a larger tibia-midfoot angle.

### Intramuscular myotenotomy

Immediately following euthanasia, the gastrocnemius muscle was exposed. A longitudinal muscle was made overlying the myotendon to expose the myotendon with care to minimally disrupt surrounding muscle and not lengthen muscle as would occur with a horizontal incision. Using microscissors, the myotendon was then transected horizontally at the level just proximal to level of myotendon pennation (Fig. 5D). Myotenotomy was performed on both right and left myotendon bands.

### Tendon composition characterization

#### Hydroxyproline

Mice were euthanized at 28DPN. Specimens (tendon or myotendon) were collected and pooled from 3-4 mice per sample and snap frozen in liquid nitrogen prior to processing. Samples were homogenized in Tissue Protein Extraction Reagent (ThermoFisher) using a tissue homogenizer as detailed in *RNA Extraction*. Collagen content was inferred by hydroxyproline quantitation by Hydroxyproline Assay Kit (Chondrex) which was performed per the manufacturer’s instructions. Pierce BCA (ThermoFisher) assays were also performed to normalize hydroxyproline values to mg of sample.

#### TEM

Whole mouse hindlimbs were fixed in 1.5% glutaraldehyde/1.5% paraformaldehyde (Electron Microscopy Sciences) in Dulbecco’s serum-free media containing 0.05% tannic acid (Sigma). Further tissue processing and transmission electron micrscopy was performed by MicroImaging Center at Shriners Children’s (Portland). Briefly, samples were then dehydrated in a graded series of ethanol to 100%, rinsed in propylene oxide, and infiltrated in Spurrs epoxy. Samples were then polymerized at 70C over 18 hours.

TEM images of transverse sections were collected at several magnifications to enable morphological visualization of the collagen fibrils and gross tendon appearance. Collagen fibril diameters were determined using ImageJ (NIH); 6 representative images were analyzed for each tendon (n = 3 independent samples per group) and a total of 2500 fibrils were analyzed per sample. Packing efficiency was assessed as total area / total summed area of collagen fibrils. Packing efficiency was determined in 3 regions of interest per tendon and averaged for each specimen. For each region of interest, total area and cumulative fibril area was quantified in ImageJ (NIH).

### Biomechanical tendon testing

For biomechanical testing, limbs were harvested at 28DPN and stored at -25C until use. On the day of biomechanical testing, limbs were thawed, and Achilles tendons were dissected out carefully with the bony calcaneal tuberosity. Mechanical tensile testing of mouse Achilles tendons was performed using custom grips to clamp the calcaneal tuberosity and Achilles tendon origin. The tendons were then immersed in a PBS bath at room temperature and preloaded to 0.05 N for ∼1min followed by a ramp to failure at 1% strain/second. Force and displacement were recorded using an Instron 8872 Universal Testing System (Instron), which was used to calculate tissue stiffness, maximum tensile force, and toughness (Golman et al., 2021).

#### *In vivo* BMP-SMAD4 inhibition

BMP-mediated Smad4 signaling was inhibited intraperitoneal administration of LDN 193189 (LDN) in sterile PBS (Tocris Bioscience). Following SNT at P5, intraperitoneal injections of LDN at 2.5mg/kg of bodyweight were performed every day from P5 till P21 (Malhotra et al., 2015). Injection site was alternated left to right to provide injection site rest. Mice injected with PBS served as controls.

### Statistics

All quantitative data are presented as means ± standard deviation with each point representing independent biological replicates. For comparison of two groups, two-tailed Student’s t-tests were performed. One-way and two-way ANOVA with Tukey’s correction for multiple hypothesis testing was performed for comparison of >2 groups. All statistical analysis was performed using GraphPad Prism v10.1.0. Sample sizes were determined based on our previous data.

## DATA AVAILABILITY

Mouse RNA-seq data is deposited in Gene Expression Omnibus GSEXXXXX.

